# Single cell transcriptomics of regulatory T cells reveals trajectories of tissue adaptation

**DOI:** 10.1101/217489

**Authors:** Ricardo J Miragaia, Tomás Gomes, Agnieszka Chomka, Laura Jardine, Angela Riedel, Ahmed N. Hegazy, Ida Lindeman, Guy Emerton, Thomas Krausgruber, Jacqueline Shields, Muzlifah Haniffa, Fiona Powrie, Sarah A. Teichmann

## Abstract

Non-lymphoid tissues (NLTs) harbour a pool of adaptive immune cells, the development and phenotype of which remains largely unexplored. Here, we used single-cell RNA-seq to characterise CD4^+^ regulatory (Treg) and memory (Tmem) T cells in mouse skin and colon, the respective draining lymph nodes and spleen. From this data, we modelled a continuous lymphoid-to-NLT trajectory for Treg, and reconstructed the mechanisms of cell migration and NLT adaption. This revealed a shared transcriptional programme of NLT priming in both skin and colon-associated lymph nodes, followed by tissue-specific adaptation. Predicted migration kinetics were validated using a melanoma-induction model, emphasizing the relevance of key regulators and receptors, including *Batf, Rora, Ccr8, Samsn1*. Finally, we profiled human blood and NLT Treg and Tmem cells, identifying cross-mammalian conserved tissue signatures. In summary, we have identified molecular signals mediating NLT Treg recruitment and tissue adaptation through the combined use of computational prediction and *in vivo* validation.

## Introduction

CD4^+^ T cells are major orchestrators of the adaptive immune response controlling host defense towards pathogens as well as maintaining tolerance towards self-antigens. In response to microbial and cytokine cues they differentiate into distinct effector cells tailored to the immune challenge and tissue location. In addition to circulating CD4^+^ T cells, recent evidence suggests that resident non-lymphoid tissue (NLT) T cell populations play an important role in providing localised immediate immune protection within the NLTs (Fan and Rudensky 2016; Watanabe et al. 2015).

Regulatory T cells (Tregs), characterized by *Foxp3* expression, are highly specialised cells which control immune responses and play a central role in homeostasis (Sakaguchi 2004; Izcue, Coombes, and Powrie 2009). Multiple studies have described unique tissue-specific adaptations of NLT Tregs distinct from their lymphoid tissue counterparts. This includes acquisition of an effector phenotype with expression of transcripts encoding effector molecules (*Ctla4*, *Gzmb*, *Klrg1*), chemokines and their receptors (*e.g. Ccr4*), and immunosuppressive cytokines such as *Il10* (*Panduro, Benoist, and Mathis 2016*; *Bollrath and Powrie 2013*). Homeostatic functions of NLT Tregs include control of insulin resistance in visceral adipose tissue (Feuerer et al. 2009), skeletal muscle repair (Burzyn et al. 2013), and hair follicle stem cell stimulation (Ali et al. 2017). Particular signature genes associated with these and other Treg populations in NLTs have been described (Liston and Gray 2014), but the full extent of their phenotype and heterogeneity within NLTs and how this changes upon challenge has yet to be uncovered. Furthermore, most studies of NLT Tregs have been performed in mice and very little is known about these cells in humans despite the enormous interest in Treg therapy and manipulation in the context of cancer, autoimmune disease, and organ transplant

Trafficking of T cells to NLTs occurs in steady-state conditions (Kimpton et al. 1995). For example, Treg were shown to be present in NLTs from a young age (Thome et al. 2015; Cipolletta et al. 2015), showing that the presence of these cells is not dependent on acute NLT immune challenges, and occurs either developmentally or in response to challenges at barrier surfaces from harmless stimuli such as commensal bacteria and dietary antigens (Ivanov et al. 2008). Migration of Tregs to NLTs requires tissue-specific cues involving cell-surface receptors (Chow, Banerjee, and Hickey 2015; Thome et al. 2015), such as chemokine receptors, as well as other G-protein coupled receptors (S. V. Kim et al. 2013) and integrins (Cepek et al. 1994).

Despite these insights into trafficking and regulation, characterization of CD4^+^ T cell populations across tissues has historically been held back by phenotypic characterization being restricted to a limited set of markers (*e.g.* single molecule FISH, FACS) or mixtures of cells (*e.g.* bulk RNA-sequencing) assayed.

To provide further insight into Treg populations in tissues we performed single-cell RNA-seq (scRNA-seq) analyses of Tregs from mouse colon, skin and compared with relevant lymphoid tissues. Our results show NLT Treg adaptations in the steady state with a core shared signature between mouse skin and colon Tregs, indicative of a general NLT residency programme in barrier tissues. Through “pseudospatial” modelling we revealed the transcriptomic adaptations occurring in Tregs during their transition from the lymph node to barrier tissues. These findings were recapitulated during *de novo* Treg recruitment to melanoma in a murine model system. Lastly, we confirmed the evolutionarily conserved and species-specific expression patterns between mouse and human Tregs. Our expression data for different tissues is also available for user-friendly interactive browsing online at data.teichlab.org.

## Results

### CD4^+^ Treg are transcriptionally distinct across tissues

We performed scRNA-seq on isolated CD4^+^Foxp3^+^ (Treg) and CD4^+^Foxp3^-^CD44^high^ memory (Tmem) T cells from two barrier NLT site: the colonic lamina propria (hereafter referred to as colon), and the skin, their lymphoid counterparts in the draining mesenteric and brachial lymph nodes (mLN and bLN) and the spleen from a Foxp3-GFP-KI mouse reporter line (Bettelli et al. 2006) (Figure 1A, Supplementary Figure 1). Cells from each NLT (and accompanying lymphoid tissues) were obtained from separate pools of mice, and henceforth we will refer to each as “Colon” or “Skin” datasets. We will refer to Treg and Tmem populations together as CD4^+^ T cells.

**Figure 1.**
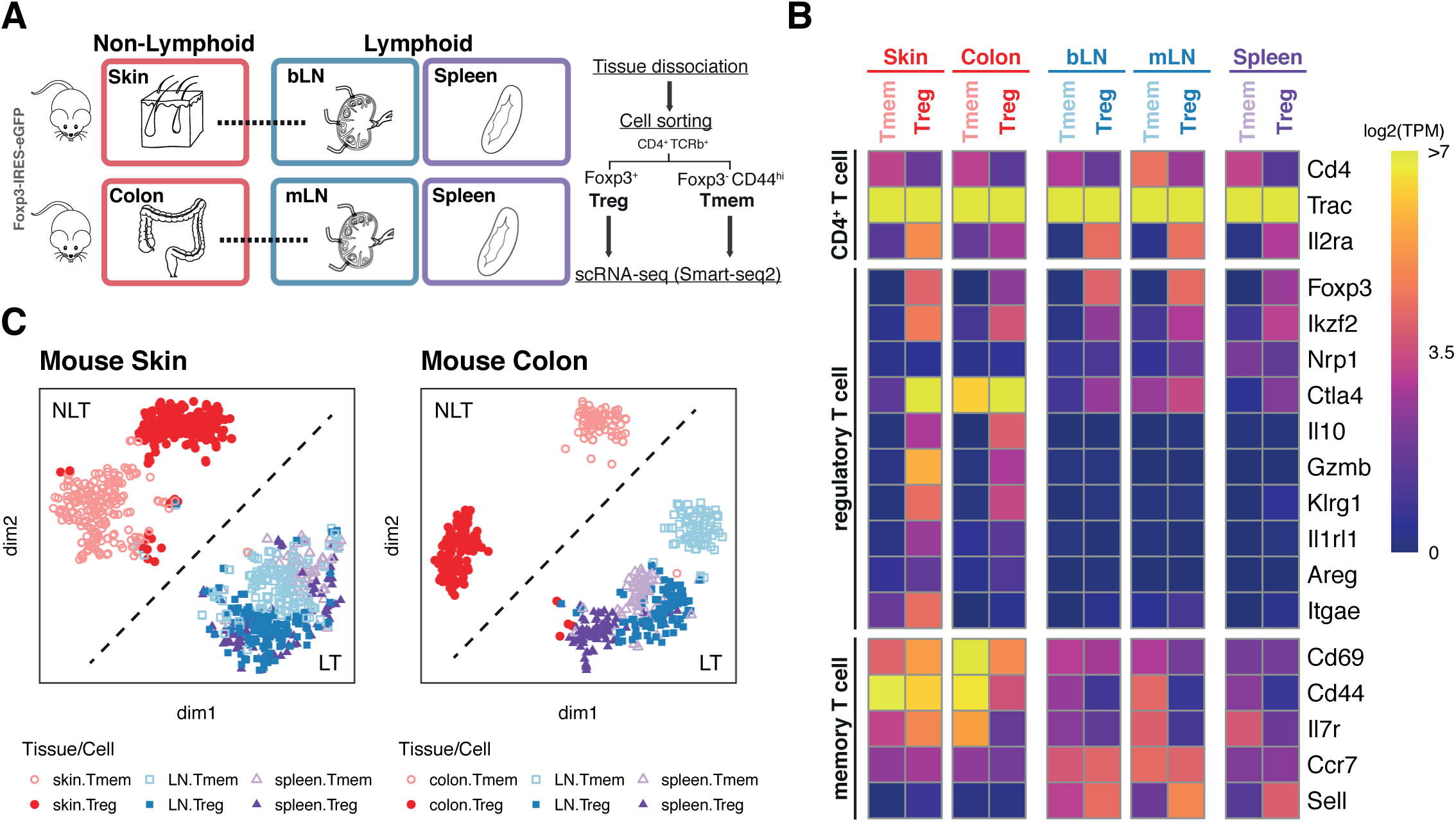
**Steady-state scRNA-seq datasets of CD4+ T cells from LT and NLT.** (A) We generated two scRNA-seq datasets composed of Treg (TCRβ^+^CD4^+^Foxp3^+^) and Tmem (TCRβ^+^CD4^+^Foxp3^−^CD44^+^) cells from an NLT (colon or skin), draining lymph node (LN, mesenteric LN for the colon and brachial LN for the skin, respectively) or spleen, sorted from Foxp3+ reporter mice.
(B) Mean expression levels of characteristic markers across CD4+ T cell populations and tissues are recapitulated by scRNA-seq.
(C and D) t-SNE dimensionality reduction of the colon (C) and skin (D) datasets, showing Treg and Tmem from NLTs (colon and skin), LNs (mLN and bLN) and spleen. Treg and Tmem are represented by filled and open symbols, respectively. Colours and symbols match tissue: NLTs in red circles, LNs in blue squares, spleen in purple triangles.

A plate-based version of the Smart-seq2 protocol was employed (Picelli et al. 2014), and after mapping and quality control (Supplementary Figure 2, see Methods) we retained 485 cells from the colon dataset, and 796 cells from the skin dataset, expressing on average 3118 genes (TPM>0) (Supplementary Figure 2).

We confirmed the expression of marker genes previously associated with Treg and Tmem in our scRNA-seq datasets (Figure 1B). *Foxp3* expression was reliably detected in Tregs and absent from Tmem cells. Effector Treg molecules such as *Ctla4*, *Il10*, *Gzmb*, *Klrg1* and *Il1rl1* (ST2), were highly expressed only in Tregs isolated from the NLTs. *Sell* (CD62L) characterised Treg and Tmem from LT (Steeber et al. 1996), while *Ccr7* is specific to lymph node Tregs, as previously described (Schneider et al. 2007).

To delineate the relationships between the T cell populations isolated, we applied tSNE to skin and colon datasets individually (Figures 1C), observing a similar cluster structure in both cases. Both skin and colonic CD4^+^ T cells were transcriptionally distinct from their respective lymphoid counterparts (Supplementary Figure 3), as previously observed in bulk RNA-seq data (Panduro, Benoist, and Mathis 2016). Treg and Tmem from spleen and LNs clustered together, albeit not overlapping, underscoring the similarity between lymphoid tissue CD4^+^ T cells.

tSNE projections did not show clear sub-clusters within each cell type in the skin or colon. To further investigate intra-population heterogeneity, we applied a consensus clustering approach to Treg and Tmem of each NLT (Kiselev et al. 2017) (Supplementary Figures 5 and 6, see Methods). This confirmed the homogeneity of colonic Tregs, with no differentially expressed genes detected between putative subclusters, in contrast to previous reports of phenotypically and functionally diverse subpopulations of CD4^+^ T cells within NLTs (Ohnmacht et al. 2015; Schiering et al. 2014; Sefik et al. 2015). While the majority of colonic Tregs express *Gata3*, the number of *Rorc*-expressing cells is low, and they do not appear to form a distinct subpopulation (Supplementary Figure 4A). Interestingly, the recently identified *Rorc*-regulator *Maf* (Wheaton, Yeh, and Ciofani 2017) is frequently expressed in both colon and skin Tregs, despite not contributing to the definition of colonic Treg subpopulations (Supplementary Figure 4B).

In contrast, the mLN Treg compartment contained two subclusters which we identified as corresponding to central Treg (cTreg, expressing *Sell*) and effector Treg (eTreg, expressing *Itgae*, *Tigit* and *Icos*) (Supplementary Figure 7D-F). Colonic Tmems also segregated into two clusters, one characterized by higher expression of Th1-associated genes, such as *Tbx21* (*T-bet*), *Il12rb* and *Ifngr1*, and another characterized by higher expression of Th17-associated genes, including *Il23r*, *Il17f* and *Il1r1* (Supplementary Figure 5). In turn, subclusters for skin Treg and Tmem differed most in metabolic activity (*Rrm1*, *Sdhd*) and cytokine signalling (*Il17f*, *Il18rap*), respectively, but did not present any clear divisions between known subtypes (Supplementary Figure 6). These findings revealed that Tregs in skin and colon were transcriptionally distinct between tissues but homogenous within one tissue.

### Gene expression signatures of NLT Treg and Tmem

Next, we investigated tissue adaptation and functional specialisations of NLT Treg. We used differential expression (DE) to determine which genes characterize CD4^+^ T cells in the NLTs. Then, to investigate cell type-specific NLT adaptations, we classified NLT markers as Treg, Tmem or shared, based on differential expression between the two populations (Figure 2A, see Methods). In colon, 36 genes were unique to Treg, 315 to Tmem, and 92 were shared between the two. In the skin, 85 genes were unique to Treg, 146 to Tmem and 527 were shared.

**Figure 2.**
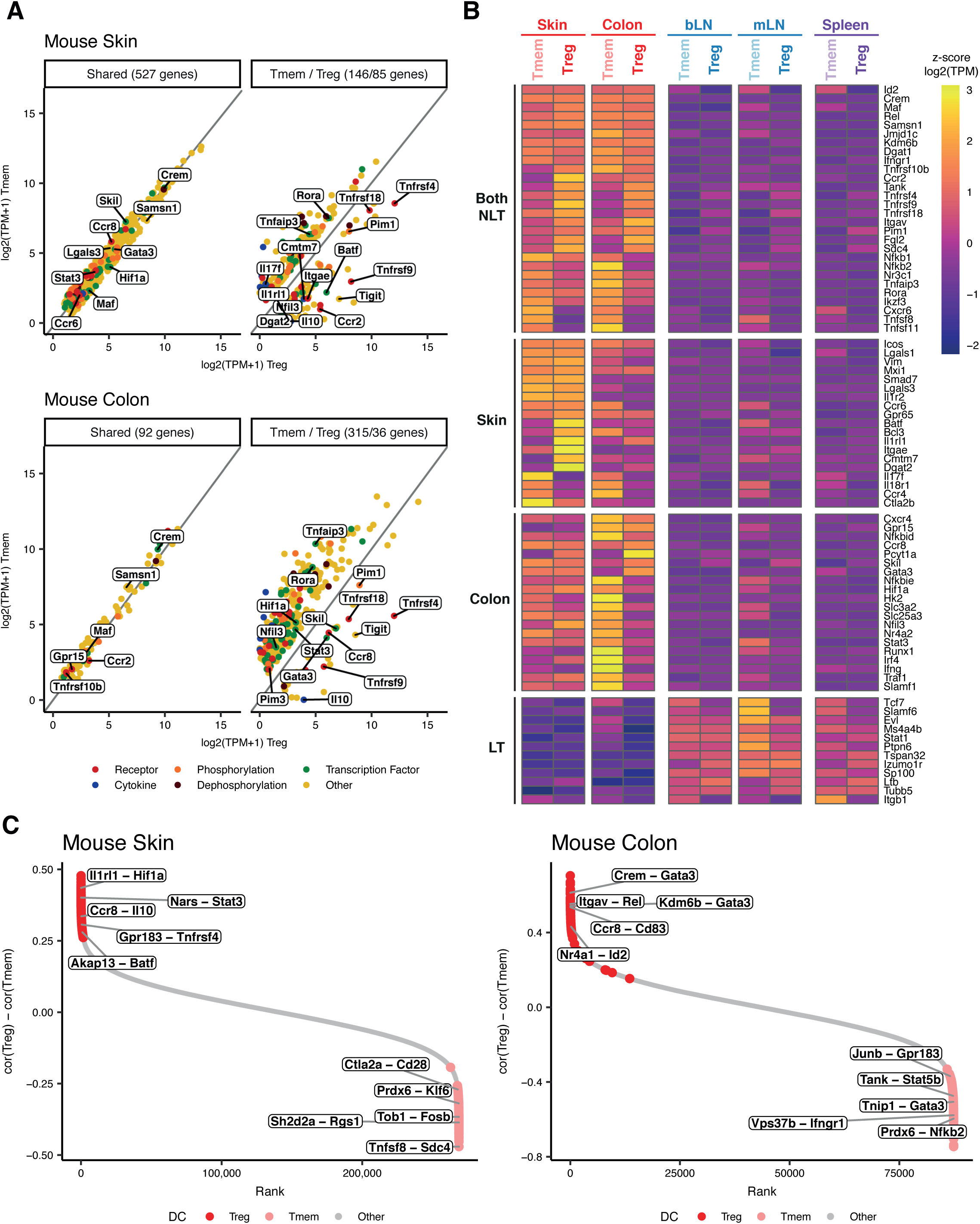
**Characterization of Treg and Tmem NLT adaptations.** A. Differential expression of NLT markers between NLT Treg and Tmem, in skin (top) and colon (bottom). Per dataset, genes were first identified as NLT- or LT-associated (q-value<=0.01, log2 fold-change (FC)>=1). This subset of genes was then characterized as Treg, Tmem or shared between cell types based on q-value<=0.05 and log2FC>=1 (see Methods).
B. Z-score of mean expression levels of newly identified markers across cell types and tissues, grouped by the populations where they are differentially expressed (left labels).
C. Differentially correlated (p-value<=0.05) genes between Treg and Tmem in mouse skin (left) and colon (right), ranked by the difference in correlation in Treg (dark red) and Tmem (light red).

NLT T cell populations are characterised by the expression of several elements of the TNFRSF-NF-*κ*B pathway, including receptors (*Tnfrsf10b*), transducers (*Traf1*, *Traf4*) and effectors (*Nfkb1*, *Nfkb2*, *Rel*). In Tmems, these are accompanied by cytokines (*Tnfsf8*, *Tnfsf11*), and various pathway inhibitors, such as *Tnfaip3* and *Tnfaip8* in both NLTs and *Nfkbid* and *Nfkbie* specifically in the colon. In contrast, NLT Tregs expressed TNF receptors (*Tnfrsf4*, *Tnfrsf9*, *Tnfrsf18*) and transducers (*Pim1*, *Tank*), underscoring the importance of signalling via the TNFRSF-NF-*κ*B axis in controlling peripheral Tregs.

Tregs and Tmems can also be distinguished by the expression of specific chemokine receptors (Figure 2A and 2B). In the colon *Ccr8* is specifically upregulated in Tregs, and *Cxcr6* is more highly expressed in Tmems. Colonic Tregs and Tmems expressed *Gpr15, Ccr2* and *Cxcr4*, as previously reported (S. V. Kim et al. 2013; J. Kim et al. 2005; Connor et al. 2004; Nguyen et al. 2014). Treg and Tmem skin populations share expression of *Ccr6* and *Gpr65*. *Ccr2* was more highly expressed in Tregs and *Cxcr6* in Tmems.

Tregs have recently been distinguished from other T helper subsets by their metabolic activity (Newton, Priyadharshini, and Turka 2016). We observed that both colon and skin Tmems express *Dgat1*, a triglyceride metabolism enzyme, whereas skin (but not colonic) Tregs express *Dgat2*. Colonic Tmems also present a higher expression of glycolysis-associated transcripts (*Hif1a*, *Hk2*), and solute carriers (*Slc3a2*, *Slc25a3*) compared to Tregs.

Other genes distinguishing these populations include regulators like *Skil* (colon Treg; expression is modulated by TGFβ (Tecalco-Cruz et al. 2012)), *Batf* (skin Treg), and *Rora* (colon and skin Tmem); extracellular matrix interactors (*Lgals1* and *Lgals3* in skin) and integrins (*Itgav* in all NLT Treg, *Itgae* only in skin Treg). We additionally found *Il17f* and *Ifngr1* present in skin Tmems, suggesting that Th1 and Th17 T helper subtypes are present in this tissue like in the colon, as expected, even though no clear subpopulations could be detected.

Most differentially expressed genes are present in both populations, albeit at different levels (Figure 2, Supplementary Figure 8). This observation suggests that individual genes may not explain all differences between populations. Therefore, we sought to identify differential gene-gene interactions in Treg and Tmem in NLTs using a differential correlation test between cell types in each NLT (see Methods). This enabled us to rank coexpressed gene pairs with cell type-specific significance. In colon (Figure 2C, right panel), *Crem* and *Gata3* are strongly correlated in Treg, mirroring their induction in Th2 cells (Sasaki et al. 2013), suggesting that regulation by *Gata3* follows similar pathways in both cell fates. Within colon Tmem, we detected a strong relationship between *Nfkb2* and *Prdx6*, which has been shown in the context of ROS homeostasis (Fatma et al. 2009) in a different system. In skin (Figure 2C, left panel), the interaction between *Il1rl1*, a known player in visceral adipose tissue (VAT) Treg development (Vasanthakumar et al. 2015), and *Hif1a*, involved in T cell metabolism reprogramming (Dimeloe et al. 2017), can be directly associated with skin Treg differentiation. Also in skin, *Stat3* potentially induces *Nars* expression (Hardee et al. 2013) in Treg, while Tmem cells exhibit signs of exhaustion by co-expressing *Sh2d2a* and *Rgs1* (Giordano et al. 2015). Overall, this method allowed us to further dissect gene interactions identified for these populations, many of them previously identified in other cells.

### Tregs share a common expression module across skin and colon

Next, we set out to directly investigate Treg adaptation in skin and colon. We extracted the unique and shared gene expression patterns across the two Treg NLT populations (Figure 3A). We identified a shared presence of *Gata3*, *Stat5a*, *Id2* and *Xbp1*, all genes involved in the regulation of T cell activation. We also found shared expression of the TNFRSF-NF-*κ*B pathway (*Tnfrsf4*, *Tank*, *Nfkbia*, *Rel*, among others) in both NLT Treg populations. Tregs from both tissues expressed other genes associated with cell survival (*Bcl2l11*, *Cdc42*), however skin Tregs expressed more genes associated with this function than colon Tregs, such as *Bcl3*, *Emp3* or *Tax1bp1*. In addition, effector Treg (eTreg) genes *Itgae, Tnfrsf9*, and VAT Treg genes *Batf* and *Il1rl1*, were upregulated in skin Tregs. Interestingly, colonic Tregs express a higher amount of immune activation and suppression genes (*Il10*, *Stat5b*, *Tigit*, *Cd83*), as well as *Erdr1*, a gene controlled by microbiota via Toll-like receptor pathways (Soto et al. 2017; Cipolletta et al. 2015).

**Figure 3.**
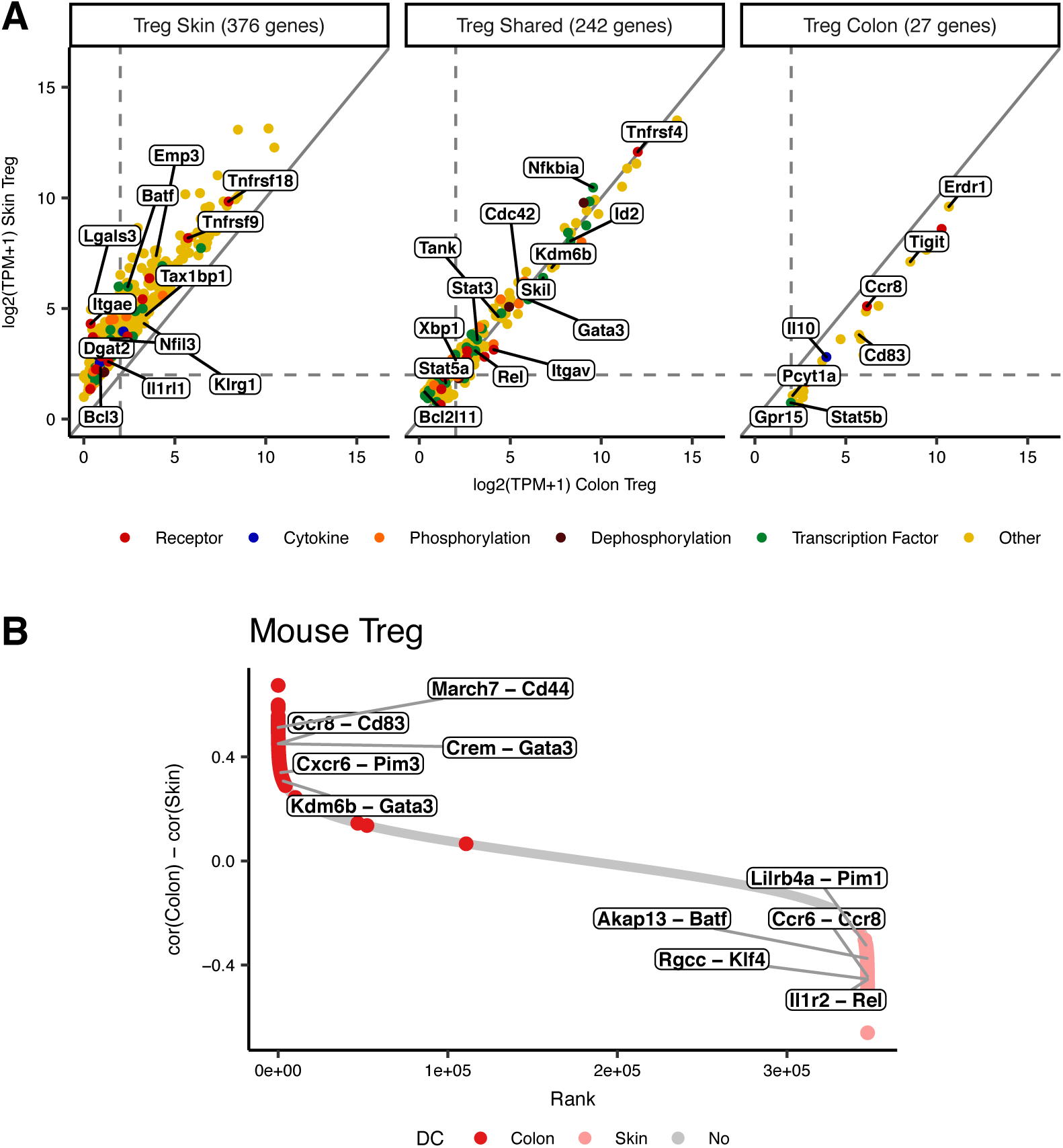
**Specific Treg adaptations to skin and colon environments.** A. Differential expression of NLT markers between skin and colon Treg. Genes are characterized as colon-specific, skin-specific or shared based on based on q-value<=0.05 and absolute log2FC>=1 (see Methods).
B. Differentially correlated (p-value<=0.05) genes between mouse colon and skin Treg, ranked by the difference in correlation in colon Treg (dark red) and skin Treg (light red).

We then examined specific gene-gene interactions of Tregs in skin and colon (Figure 3B). The differential correlation test highlighted several pairs of genes which shift in relevance from colon to skin. For example, in colon Treg, previously described MARCH-mediated ubiquitination effects on *Cd44* stability (Nakamura 2011) might underlie *March7* correlation with *Cd44*. In skin, we observe a high correlation between *Rel* and *Il1r2*, whose promoter contains a potential binding site for *Rel* (Gagné-Ouellet et al. 2015), and a high correlation between the MAPK-pathway genes *Rgcc* and *Klf4*, which have been shown to be correlated in a T cell context (Michel et al. 2017).

### Tissue adaptation dynamics of NLT Tregs

It is well recognized that Treg primed in LT are subsequently recruited to NLT (Agace 2006). However, the mechanisms underlying recruitment of Tregs from LT to NLT is far from being understood.

First, to obtain evidence of CD4^+^ T cell recruitment from LT to NLT in our data, we checked for TCR clonotypes shared between tissues using TraCeR (Stubbington et al. 2016), as a strategy to fate map Treg clones across lymphoid and non-lymphoid tissues. We identify Tmem and Treg clones present in both lymph nodes and NLTs (Supplementary Figure 9), suggesting migration between them. Additionally, TCR clonotype analysis confirmed that clonal relationships occur exclusively within the Tmem and Treg compartments, rather than between them, suggesting that in these tissues under homeostatic conditions Tregs and Tmems do not exhibit a precursor product relationship.

We then hypothesised that gradual transcriptional changes within Tregs are likely to represent their adaptation to the respective NLT environments. Furthermore, the identification of an effector-like subset in mLN Treg with increased potential to migrate out of lymphoid tissues (suggested by the downregulation of *Sell* (CD62L); Supplementary Figure 7), links tissue adaptation and migration from LN to NLT.

To identify these trends in the data, we computationally reconstructed a pseudospace relationship between cells. We employed Bayesian Gaussian Process Latent Variable Modelling (BGPLVM, see Methods) (Michalis K Titsias 2010) to Treg transcriptomes from NLTs and their respective draining-LN. The non-linear latent variables (LVs) order cells according to the gene expression patterns they represent, and can then be interpreted in this biological context. For each dataset, we identified two main LVs that explain most of the transcriptomic differences present across cells and tissues (LV0 and LV1) (Supplementary Figure 10A,B). Importantly, the general cell distribution from each tissue along LVs (Figure 4A), as well as the genes associated with each of them are similar in both mLN-to-colon and bLN-to-skin transitions (Supplementary Figure 10C; see Methods). We also observe that this pair of latent variables contains a larger proportion of NLT-associated genes (Supplementary Figure 10D), and the gradient is retained even when analysing individual tissue populations (Supplementary Figure 11). Thus, as a result of the demonstrated association to NLTs marker genes, LV1 in colon and LV0 in skin can be used as a pseudospace variable to represent the continuum of Treg trafficking and adaptation between LNs and NLTs.

**Figure 4.**
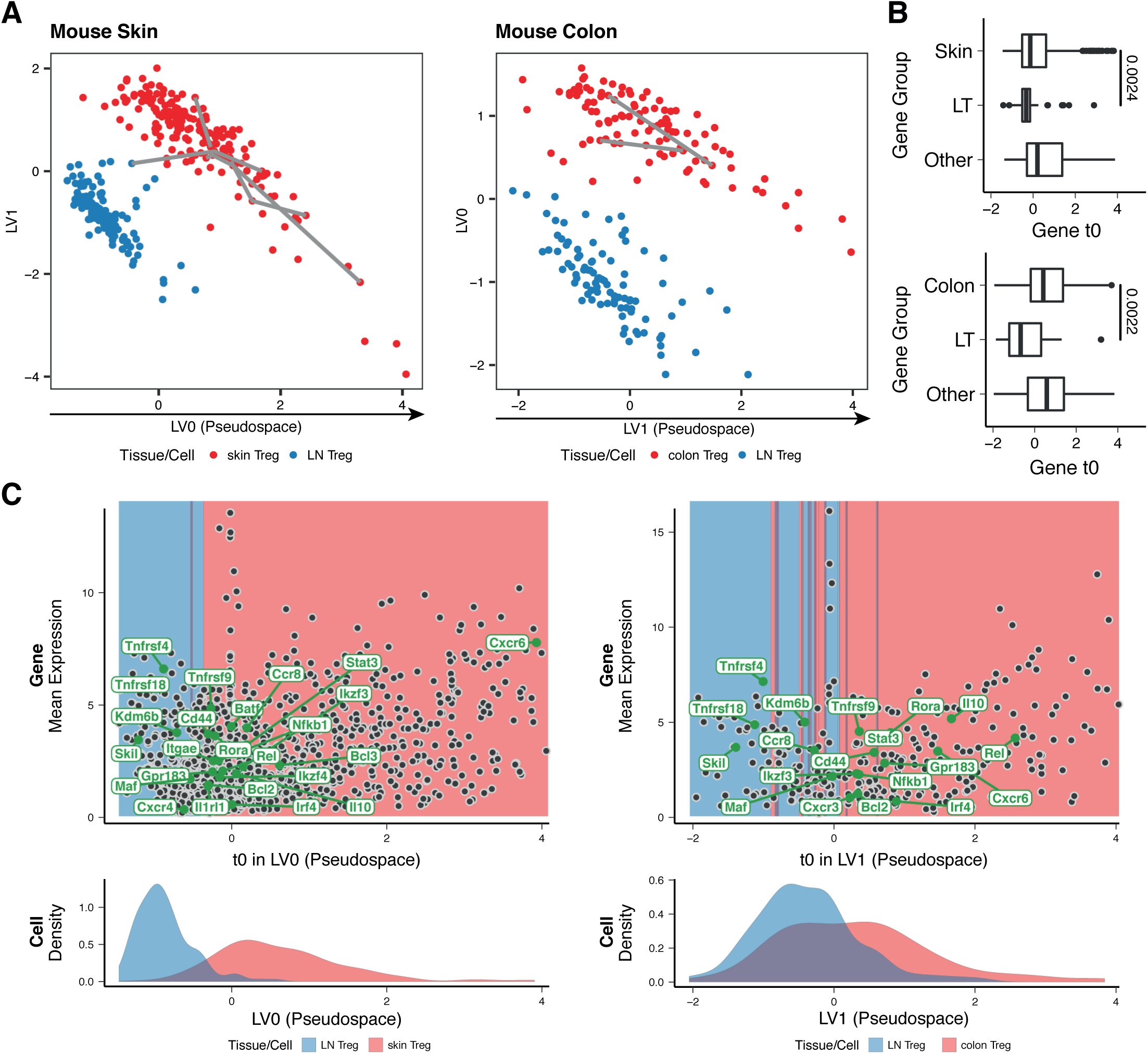
**Reconstruction of Treg recruitment from lymphoid to non-lymphoid tissues in steady-state.** A. Top two latent variables found with BGPLVM of Treg in skin (left) and colon (right) datasets (see Supplementary Figure 11), associated with either tissue of origin or activation/recruitment to non-lymphoid tissue. Colours match tissue of origin: non-lymphoid tissue in red, lymph nodes in blue. Grey lines connect cells sharing the same productive α and β TCR chains, mentioned in the main text as clonotypes.
B. Distribution of activation/deactivation points (t0) for NLT (top - skin; bottom - colon) and lymphoid tissues (LT) markers, and remaining genes.
C. Activation/deactivation point (t0) and mean expression (μ0) of genes significant in migration to colon or skin. (Top) Point of activation/deactivation (t0) of genes along the activation/recruitment trajectory as determined by the switchDE package, in skin (left) and colon (right) datasets. (Bottom) Distribution of Treg cells from lymph nodes and the non-lymphoid tissues along trajectories.

Having established a pseudospatial trajectory for recruitment and adaptation of Tregs from LNs to NLTs, we next examined the kinetics of up- and down-regulation of genes along it, with the goal of identifying the cascade of transcriptional changes that ultimately constitute the adaptation to non-lymphoid tissues. We addressed this question by fitting a sigmoid curve to the expression values of each gene along this trajectory (Supplementary Figure 12, see Methods). This allowed us to determine that 1163 and 318 genes have a switch in expression (either up or down) along the bLN-to-skin and mLN-to-colon paths, respectively, with 243 genes in common between the two trajectories. These are enriched in cell activation, adhesion and localization functions (Supplementary Figure 14), cell processes fundamental for activation and migration of immune cells, and have a high representation of NF-*κ*B and TNF signaling pathways linked to eTreg differentiation and maintenance (Grinberg-Bleyer et al. 2017; Vasanthakumar et al. 2017). The kinetics of these core genes are particularly conserved in the initial stages of Treg NLT adaptation, being enriched in “leukocyte transendothelial migration” and “cytokine-cytokine receptor interaction” KEGG pathways (Supplementary Figure 15). Later stages of gene expression changes are less consistent between colon and skin trajectories, which can potentially be explained by different environmental cues in each NLT. Nevertheless, the transcription factors *Stat3* and *Irf4*, known to be important in Treg adaptation and differentiation (Chaudhry et al. 2009; Cretney et al. 2011), are part of the final wave of a shared adaptation signature.

The point of activation/deactivation of each gene in the trajectory (t0) (Figures 4C) shows that genes previously identified as markers for non-lymphoid tissues have a significantly later activation time than genes from lymphoid tissues (Figure 4B), in line with the tissue distributions presented. Nonetheless, some NLT Treg-associated genes appear to be upregulated already in the lymph nodes.

Based on cell neighbourhood, we then divided the pseudospace into LN and NLT regions (Figure 4C), to identify where in the trajectory gene expression changes take place - in the lymph node earlier in the trajectory, or in the NLT itself. TNFRSF-NF-*κ*B-related genes (*Tnfrsf4*, *Tnfrsf18*) are upregulated in the LN, reflecting the presence of eTreg traits in the lymphoid compartment, and the relevance of this signalling pathway for the development of the NLT phenotype. The upregulation of the transcription factor *Skil* and the histone demethylase *Kdm6b* in cells in the lymph node, and continuous expression in the NLT cells, points towards a role for these genes in the initial stages of tissue adaptation. The NLT section of the pseudotime shows evidence of further Treg differentiation, containing additional players involved in the NF-*κ*B pathway (*Rel, Nfkb1, Tnfrsf9*), and upregulation of effector molecules (*Il10*, *Cd44*). Important regulators for the final tissue adaptation include *Rora*, *Ikzf3*, *Irf4*, and *Stat3*. Simultaneously, the need to retain Tregs in NLTs is met by the upregulation of membrane receptor molecules: *Ccr8*, *Gpr183*, *Cxcr6*. *Cxcr4* and *Cxcr3* are expressed in the skin and colon trajectories respectively, suggesting they should differentially contribute to localisation in distinct NLTs. Skin cells express additional markers that indicate increased survival ability (*Bcl3*) and further adaptation to non-lymphoid environments (*Batf*, *Il1rl1*, *Itgae*, *Ikzf4*).

In summary, we were able to capture a range of Treg stages and order cells across lymphoid and non-lymphoid tissues, establishing a continuous trajectory of recruitment and adaptation. Furthermore, from their conserved sequential order of activation, we conclude that gene activation leading to NLT adaptation follows a similar regulatory sequence in both bLN-to-skin and mLN-to-colon trajectories, with a core set of shared gene expression patterns reflecting a strong influence of the TNFRSF-NF-*κ*B axis.

### Treg recruitment into steady-state skin and melanoma tumours utilises shared mechanisms

We dissected the relationship between lymph node and NLT Tregs in the steady-state, by providing evidence for a tissue adaptation trajectory using transcriptomic similarity and clonal relationships. To validate our findings, we now focus on a perturbed system where Treg recruitment is induced to see whether we can recapitulate their migration and adaptation trajectory to NLTs. Tregs are known to be recruited into tumours, where they contribute to the immunosuppressive environment that facilitates tumour growth. Importantly, previous studies analyzing human TCR repertoires (Sherwood et al. 2013; Plitas et al. 2016) have shown that tumour-Treg are likely to be recruited *de novo* from lymphoid tissues and not from the adjacent NLT, despite exhibiting a phenotype similar to that of NLT Treg (Plitas et al. 2016; De Simone et al. 2016). Thus, we purified NLT CD4^+^ T cells from B16.F10 melanomas or PBS controls 11 days after subcutaneous implantation into Foxp3-IRES-eGFP reporter mice (Haribhai et al. 2007) (Figure 5A, Supplementary Figure 16; see methods).

**Figure 5.**
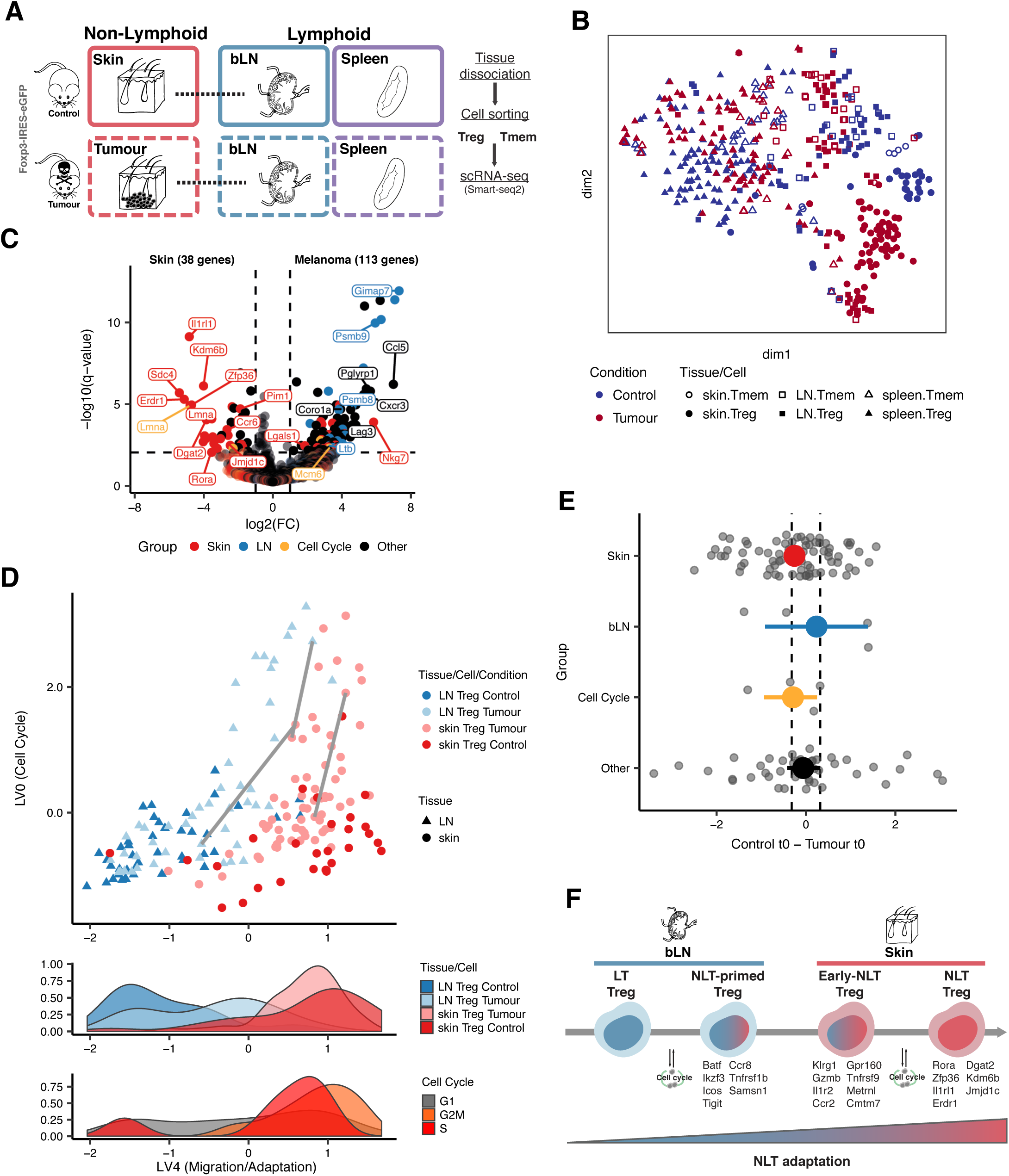
**Recruitment and adaptation of Treg to the tumour environment recapitulates steady-state migration.** A. Melanoma induction strategy and tissues sampled for different cell-types (similar to Figure 1A, see Methods).
B. t-SNE dimensionality reduction of single-cell data from control and tumour conditions, depicting Treg and Tmem steady-state skin and tumour, draining brachial lymph nodes and spleen. Treg and Tmem are represented by filled and open symbols, respectively. Colours match condition: control in dark blue, melanoma in dark red. Symbols match tissue: non-lymphoid tissues in circles, lymph nodes in squares, spleen in triangles.
C. Differential expression between skin and tumour (non-cycling) Treg (q-value<0.01 and log2(FC)>1).
D. (top) Latent variables found with BGPLVM representing cell cycle and non-lymphoid tissue recruitment/adaptation of Treg (Supplementary Figure 18, see Methods). Tissue of origin matches colour: skin in red, lymph node in blue. Condition matches shade: control cells darker, tumour cells lighter. Grey lines connect cells sharing the same productive α and β TCR chains. (bottom) Distribution of cells in different categories (Tissue and Condition, Cell Cycle phase) along the recruitment trajectory.
E. Comparison between time of activation of genes (t0) in control and tumour, measured as the difference of t0 between conditions. Genes are classified as being markers of skin, lymph node, cell cycle or other. Coloured points show mean +/- mean standard error. Vertical dashed lines represent +/- difference between mean t0 in each condition.
F. Proposed model of gene expression changes in NLT adaptation of Tregs. At each stage examples of genes activated are listed below.

As seen in the steady-state skin dataset, we observe shared clonotypes between tumour Tregs and bLN Tregs (Supplementary Figure 9). In the tumour-bearing mice, we detected an additional cluster of cycling cells in both the LN and tumour (Supplementary Figure 17). Cycling Tregs in tumours and respective draining-LN, combined with clonotype sharing between both locations, suggest de novo Treg recruitment from LN and simultaneous expansion in both tumour and draining-LN.

Skin and tumour Tregs clustered separately with tSNE (Figure 5B). Differential expression analysis between non-cycling tumour Tregs and control skin Tregs reveals a relatively small number of genes are significantly different between the two populations of Treg (112 upregulated in tumour Treg and 37 in steady-state skin Treg (Figure 5C)), in line with recently published human data (Plitas et al. 2016). Unsurprisingly, tumour Tregs upregulate genes related to proteasome activity, along with other pathways indicative of immune activation (Supplementary Figure 18). They also upregulate the exhaustion marker *Lag3* (Malik et al. 2017), as well as *Cxcr3*, *Nkg7*, *Lgals1*, *Ccl5,* and some LN markers, such as *Ltb*. Control skin Tregs, on the other hand, show upregulation of skin Treg markers such as *Il1rl1*, *Rora*, *Pim1*, *Sdc4*, *Kdm6b*, *Jmjd1c* and *Erdr1*. Despite these differences, tumour and skin Treg are overall very similar in terms of recruitment and NLT adaptation. Skin Treg signature genes such as *Batf*, *Tnfrsf4*, *Tnfrsf9*, *Samsn1*, *Tigit*, *Tchp*, *Ccr8*, *Ccr2* and *Itgav* are expressed at equivalent levels in skin and tumour Treg.

Next, we sought to obtain a shared migration trajectory of steady state vs perturbed system (tumour model) Treg NLT recruitment. To this end, we used the MRD-BGPLVM algorithm (Andreas Damianou, Carl Ek, Michalis Titsias, Neil Lawrence 2012) (see Methods) to explore gene expression trends across Treg from the control skin, tumour and respective draining-LNs together. Two main latent variables were identified, one concentrating most of cell cycle-associated variability (LV0), and one mainly associated with the NLT signature (LV4) (Supplementary Figure 18). Notably, NLT adaptation trajectory (LV4) correlates with trajectories found in control and melanoma conditions when MRD-BGPLVM is applied to each one individually (respectively, 0.63 and 0.45 Spearman correlation coefficients; Supplementary Figure 19A-D, see Methods). Moreover, it also correlates with the NLT adaptation pseudotime previously determined for steady-state skin (0.35 Spearman correlation coefficient, Supplementary Figure 19E). These observations support the idea that changes in Treg gene expression leading to NLT adaptation along bLN-to-skin and bLN-to-tumour trajectories are equivalent processes.

We investigated gene kinetics along NLT adaptation (LV4) separately in control and melanoma conditions. 129 genes are shared between both conditions, 73% of which were also present in the steady-state skin trajectory determined previously. As we expected, values of t0 remain largely unchanged between control and melanoma (Figure 5E), further suggesting that NLT recruitment and adaptation follow the same program in homeostatic and perturbed conditions. The tissue adaptation genes shared between control and melanoma include many of the players in the TNFRSF-NF-*κ*B we previously described in the steady-state (*Tnfrsf9*, *Tnfrsf4*, *Tnfrsf18*, *Bcl3*). These are accompanied by genes encoding for proteins associated with cell migration and adhesion (*Ccr2*, *Ccr8*, *Itgav*, *Plxna2*, *Iqgap1*), secreted factors (*Lgasl1*, *Metrnl*), and others related to immune activation and effector states (*Il1r2*, *Klrg1*, *Icos*, *Tigit*, *Ctla4*). We additionally confirm the relevance of a variety of transcription factors (*Rora*, *Ikzf2*, *Ikzf3*, *Id2*, *Batf*, *Nfil3*) for migration and adaptation to the skin (Supplementary Figure 20).

Despite the similarities between melanoma and control, the distributions of cells from both conditions are not completely overlapping along the NLT adaptation trajectory, and can be ordered by NLT adaptation between populations (from least to most adapted: control LN Treg, melanoma LN Treg tumour Treg and, control skin Treg) (Figure 5D). This implies that in response to an ongoing challenge in the peripheral tissue, a higher fraction of Tregs in the LNs acquires NLT adaptations. In fact, for several NLT markers we do observe more cells expressing them (TPM>0) in the tumour-draining LN compared to the control, *e.g. Id2* (53% vs 24%), *Batf* (49% vs 17%), *Nfil3* (43% vs 21%), *Lgals1* (85% vs 57%), *Ccr8* (72% vs 45%), *Samsn1* (83% vs 55%). This further supports our claim that there is priming of Treg to NLTs while still in the LN. Finally, and as suggested by DE analysis (Figure 5C), tumour Tregs have the most cells with NLT adaptation, which brings them very close to the control skin Tregs. Overall, Tregs from challenged mice recapitulate and fill in the gaps along steady-state NLT adaptation. We summarise this trajectory in Figure 5F, where we show the stages of adaptation and key markers for each stage.

### Evolutionary conservation between mouse and human NLT Treg cells

To address how translatable our findings are to human biology, we complemented our extensive characterization of murine NLT CD4^+^ Treg and Tmem by comparing to their human counterparts. We collected Treg, Tcm and Tem cells from blood and skin of individuals undergoing mammary reduction surgical interventions, and also from tumour-adjacent colon sections from patients undergoing colonic resection (Figure 6A, Supplementary Figure 21). Similar to the mouse analysis, we identified gene markers for human CD4^+^ T cell populations (Supplementary Figure 22, see methods).

**Figure 6.**
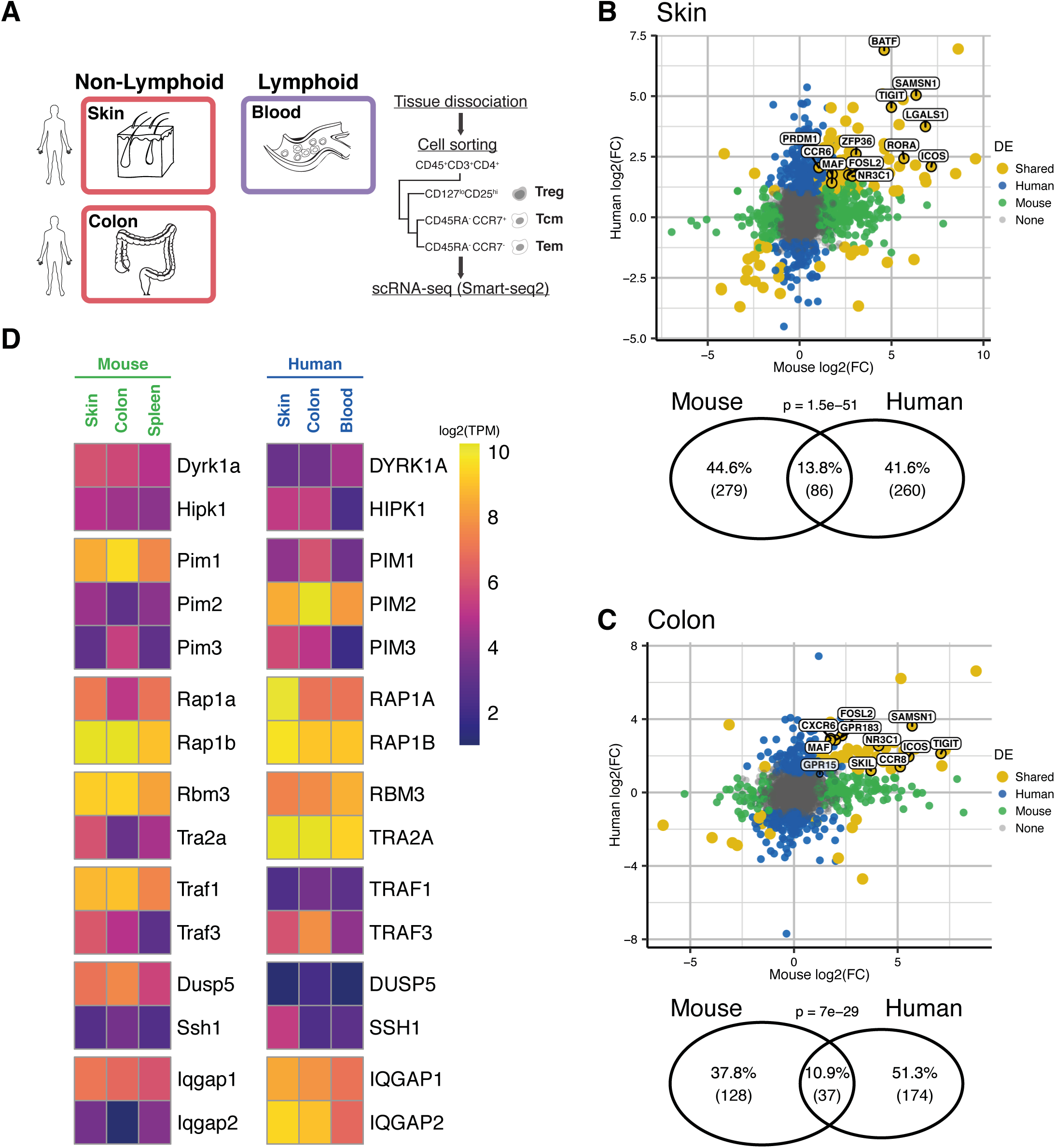
**Human-mouse comparison of non-lymphoid tissue Treg marker genes.** (A) Tissues sampled for different cell-types from human individuals (similar to Figure 1A, see Methods).
(B and C) Top: Overlap between non-lymphoid tissue Treg markers detected in human and mouse, in either (B) colon or (D) skin datasets. Bottom: Fold-change between gene expression in non-lymphoid and lymphoid tissues in mouse and human. Blood and spleen were used as lymphoid tissues in human and mouse respectively.
(D) Non-lymphoid tissue paralogs exhibiting opposing expression patterns between human and mouse.

Focusing on one-to-one orthologs, we found that 86 out of 350 human skin Treg markers and 37 out of 214 human colon Treg markers overlap with the respective mouse signature, a statistically significant level of conservation. The general NLT signature common to both organs includes human/mouse conserved TNFRSF-NF-*κ*B-pathway receptors (*Tnfrsf1b*, *Tnfrsf4*, *Tnfrsf18*) and a regulator (*Tank*), T cell activation markers (*Ctla4*, *Icos*, and *Cd44*), and transcriptions factors (*Maf* and *Fosl2*). In tissue-specific terms, *Rora*, *Prdm1*, *Batf*, *Zfp36*, *Ccr6* and *Lgals1* in skin Treg, and *Skil*, *Ccr8*, *Cxcr6* and *Gpr183* in colon Treg are markers conserved across both organisms (Figure 6B and 6C). Most of these genes were also detected in the pseudospatial trajectories obtained from the mouse data, hinting at conserved migration and adaptation mechanisms across mammals.

Close inspection of NLT markers obtained for both species revealed several instances where the expression pattern of one gene is substituted by a paralog in the other organism (Figure 6D). For example, while the kinase *Pim1* is a marker of mouse NLT Tregs, and is not expressed in human, the converse is true of *Pim3*. A similar situation was observed for *Ccr2*/*Ccr4*, *Traf1*/*Traf3* and others. This suggests that some paralogous proteins have evolved to substitute for each other during the evolution of NLT Tregs in mammals. The fact that several of the identified cases are receptors of related to signal transduction leads us to believe that evolution of cell-cell communication pathways owes some plasticity to differential paralog usage.

Our cross-species comparison suggests that despite mouse-human differences, the NLT Treg adaptation program defined in mouse is generally conserved in human.

## Discussion

It has become increasingly apparent that NLT Treg populations contribute to immune and non-immune functions. Phenotypic differences between NLT and LT populations support tissue-specific functions and enable NLT CD4^+^ T cells to overcome the challenges presented by the non-lymphoid environment. Understanding NLT adaptations is therefore of fundamental biological interest, and can ultimately help harness NLT CD4^+^ T cells for future therapeutic applications.

Our work sheds light on the phenotype of skin and colon CD4^+^ T cells. By profiling both NLT Treg and NLT Tmem, we were able to identify global relationships between cell populations, discriminating between markers shared across NLT CD4^+^ T cells and those specific to Tregs. Importantly, detected lineage-specific markers in steady-state may be distinct from previously reported studies on immune challenged tissues, even though we show a strong similarity between Treg trafficking in steady-state and melanoma (Figure 5E). We found that these Treg populations present fundamental traits shared across the skin and colon compartments, namely a substantial prevalence of genes part of the TNFRSF-NF-*κ*B axis. We verified that skin Treg adopt a phenotype closer to a VAT-Treg signature (*Itgae, Batf* and *Il1rl1*) (Hu and Zhao 2015), while colonic Treg are in a more activated and suppressive state (*Il10*, *Stat5b*, *Tigit*, *Cd83*). Interestingly, these differences do not seem to be associated with different origins of Tregs (tTreg vs pTreg), as tTreg markers *Gata3* and *Ikzf2* (HELIOS) are expressed by Tregs in both NLTs, while the pTreg marker *Rorc* (RORγt) is nearly absent from all Treg populations, but still detected in colonic Tmems (Supplementary Figure 4B, bottom).

Despite these differences between NLTs, Treg and Tmem populations within each tissue are surprisingly homogeneous, with the exception of colonic Tmems, which displayed subpopulations driven by Th1- and Th17-associated genes (Supplementary Figures 5 and 6). The absence of the previously identified RORγt^+^ and T-Bet^+^ (*Tbx21*) Treg populations (Koch et al. 2009) is likely related to the lack of exposure of the mice to significant immune challenges. In fact, we see a small proportion (20%) of RORγt^+^ colonic Tregs by intracellular staining (Supplementary Figure 4A), suggesting a limited challenge by gut microbiota in our facility. On a related note, as expected, while the protein-RNA correlations on a per-cell basis are not high, protein and RNA average values across cell populations are uniformly highly correlated (Supplementary Figure 3C).

We identified the core signature for adaptation of CD4^+^ T cells to the non-lymphoid environment. On top of this, additional features are added to attain the complete skin and colon-specific phenotypes. This modular nature of the NLT identity is evident from cell surface protein expression: *Itgav* and *Gpr183* are part of a general NLT Treg signature, and appear to set the stage for migration and retention in NLTs, while a second level of receptors, *e.g. Gpr15* (S. V. Kim et al. 2013) in colon and *Ccr6* (Scharschmidt et al. 2017) in skin, then determines tissue-specific Treg retention. These classes of genes are especially relevant for translational applications in cell-based Treg therapies, and can potentially be used to “address” cells to NLT locations. Indeed, we observe an expression of *Gpr15* and *Ccr6* in human colon and skin, respectively, which points towards a conserved mechanism of tissue specification for Treg. Future studies on more human tissues will further unravel these tissue specification mechanisms.

Understanding phenotypic variation goes beyond identifying subpopulations of cells. Having established the tissue-specific differences and similarities between regulatory T cell populations, we used Bayesian Gaussian Process latent variable modeling to extract pseudospatial relationships between cells purified from the draining lymph nodes and NLTs. Based on this alignment of cells, we were able to explore the basic mechanisms for the establishment of peripheral Treg phenotypes. The transcriptional changes associated with this transition from a continuous trajectory that starts in the draining-lymph nodes and continues in the NLTs, both in steady-state and immune challenge conditions. In the first case, the order of transcriptional activation and repression for core genes is similar between skin and colon, particularly in the first half of the trajectory (Supplementary Figure 14 and 15), pointing at a conserved programme for cell trafficking between LN and NLTs. The patterns and kinetics of TNF receptors (*Tnfrsf4*, *Tnfrsf9*, *Tnfrsf18*) and additional players in NF-*κ*B signaling highlight the role that this pathway exerts over Treg function and survival in the NLTs as a whole, both in mouse and in human, in line with recent reports (Vasanthakumar et al. 2017; Grinberg-Bleyer et al. 2017).

We increased our resolution of the expression space during Treg transitioning from lymphoid to non-lymphoid tissues by using a tumour-induction model. Here, the presence of cycling cell populations in both the tumour and the draining-LN, in conjunction with clonotype-sharing between tumour and LN Tregs, suggests *de novo* recruitment of Tregs to NLTs, in line with recent reports on human tumour-infiltrating Tregs (Plitas et al. 2016). Interestingly, a considerable proportion of the adaptation programme between bLN-to-tumour is contained within the bLN-to-skin and mLN-to-colon trajectories (Supplementary Figure 18B). Cues derived from NLTs are likely to trigger transcriptomic changes in Tregs located in the draining-LNs, placing them closer to the NLT-adapted phenotype, as indicated by a higher percentage of cells expressing *Batf*, *Ccr8*, *Samsn1* and other NLT markers in the melanoma condition. However, and despite the overall minor differences between control and melanoma conditions, the NLT trajectory suggests that tumour Tregs are less mature versions of homeostatic skin Tregs.

The trajectory of LT-NLT trafficking is supported by Treg clonotype sharing between tissues, which we infer by reconstructing the TCR α and β chains from each cell’s full-length mRNA transcriptome. The expanded clonotypes are exclusively found within Tregs or Tmems for both species (Supplementary Figure 9). In other words, Treg and Tmem are generally distinct and committed cell fates *in vivo* rather than continuously interchangeable states, in agreement with recent work by Bacher and colleagues (Bacher et al. 2016).

While we find many parallels between mouse and human Treg populations across multiple levels, there are exceptions. Specific paralogues present in both species, in particular in the PIM and the TRAF families (Figure 6F), and they are deployed differentially in each of the two species for the same function. This suggests a pivotal role for expanded gene families in rewiring signalling pathways throughout evolution. Integrative studies focusing not only on tissue-resident cells, but also on the surrounding environment and organs can help dissect the relevance of these pathways in T cell biology, and how this evolutionary rewiring might affect immune response and homeostasis.

Overall, our results reveal a dynamic adaptation of T cells as they traffic from one tissue to another, and provide an open resource (available at data.teichlab.org) for investigating *in vivo* CD4^+^ T cell phenotypes in mouse and human.

## Methods

### Mice

All mice were maintained under specific pathogen-free conditions at the Wellcome Trust Genome Campus Research Support Facility (Cambridge, UK) and at the Kennedy Institute for Rheumatology (Oxford, UK). All procedures were in accordance with the Animals Scientific Procedures Act 1986. For steady-state experiments, the Foxp3-GFP-KI mouse reporter line (Bettelli et al. 2006) was used. The melanoma challenge was performed in Foxp3-IRES-GFP knock-in reporter mice (Haribhai et al. 2007) purchased from The Jackson Laboratory (stock no. 006772). In both cases, 6-14 week-old females were used.

### Human samples

Human skin and blood samples were obtained from patients undergoing breast reduction plastic surgeries (REC approval number: 08/H0906/95+5).

Surgical-resection specimens were obtained from patients attending the John Radcliffe Hospital Gastroenterology Unit (Oxford, UK). These specimens were obtained from normal regions of bowel adjacent to resected colorectal tumours from patients undergoing surgery. Informed, written consent was obtained from all donors. Human experimental protocols were approved by the NHS Research Ethics System (Reference number:11/YH/0020). Further details concerning patients and tumours can be found in Supplementary Table 3.

#### Isolation of murine leukocytes for steady-state skin dataset

To isolate leukocytes from ear tissue, ears were removed at the base, split into halves and cut into very small pieces. Tissue was digested in 3.5ml RPMI media with 0.1% BSA, 15mM Hepes, 1mg/ml collagenase D (Roche) and 450μg/ml Liberase TL (Roche) for 60 minutes at 37°C in a shaking incubator at 200rpm. Digested tissue was passed through an 18G needle to further disrupt the tissue and release cells. Cells were passed through a 70μm cell strainer, and the digestion was terminated by addition of ice-cold RPMI containing 0.1%BSA and 5mM EDTA. A three-layer (30/40/70%) Percoll density-gradient was used to enrich for the lymphocytes. Cells obtained from the digestion were layered in the 30% layer on top of the 40% and 70% layers, and centrifuged for 20 minutes at 1800rpm without brake. Cells at the 40/70% interface were collected for the subsequent analysis. Cell suspensions from spleen and bLN were prepared as described previously (Uhlig et al. 2006).

#### Isolation of murine leukocytes for steady-state colon dataset

Colons were washed twice in RPMI media with 0.1%BSA and 5mM EDTA in a shaking incubator at 200rpm at 37°C to remove epithelial cells. The tissue was then digested for an hour in the presence of RPMI/10%FCS/15mM Hepes with 100U/ml collagenase VIII. Digestion was terminated by addition of ice-cold RPMI/10% FCS/5mM EDTA. A three-layer (30/40/70%) Percoll density-gradient was used to enrich for the lymphocytes. Cells obtained from the digestion were layered in the 30% layer on top of the 40% and 70% layers, and centrifuged for 20 minutes at 1800rpm without brake. Cells at the 40/70% interface were collected for the subsequent analysis. Cell suspensions from spleen and mLN were prepared as described previously (Uhlig et al. 2006).

### Melanoma induction and cell isolation

The melanoma induction experiments were performed in accordance with UK Home Office regulations under Project License PPL 80/2574. Protocol used was adapted from a previous publication (Riedel et al. 2016). For syngeneic tumours, 2.5 × 10^5^ B16.F10 melanoma cells were inoculated subcutaneously into the shoulder region 6- to 14-week-old female Foxp3-IRES-GFP mice (Haribhai et al. 2007). Animals were excluded only if tumours failed to form or if health concerns were reported. Control Foxp3-IRES-GFP mice were injected with 50 μl PBS. Animals were culled after 11 days. Tumour tissue, tumour-draining (brachial) lymph nodes and spleen were isolated for subsequent analysis. PBS-injected and steady-state skin, draining lymph nodes (bLN) and spleen were collected from control mice. Tumour and PBS-injected skin were mechanically disrupted and digested in a 1ml mixture of 1 mg/ml collagenase A (Roche) and 0.4 mg/ml DNase I (Roche) in PBS (solution A) at 37°C for 1h with 600rpm rotation. 1ml of PBS containing 1mg/ml Collagenase D (Roche) and 0.4 mg/ml DNase I (solution B) was then added to each sample, which returned to 37 °C for 1h with 600 rpm rotation. Lymph nodes were digested for 30min in 500μl of solution A, and for further 30min after the addition of 500μl of solution B. EDTA at the final concentration of 10mM was added to all samples. Spleens were processed as described previously (Uhlig et al. 2006). Suspensions were passed through a 70 µm mesh before immunostaining with combinations of fluorescently conjugated antibodies (Supplementary Table 1). Samples from different animals were kept separated throughout processing and sorting. Cells from PBS-injected and steady-state skin were compared and concluded to be equivalent in expression (Supplementary Figure 16E).

### Isolation of human CD4+ T cells

#### Isolation of leukocytes from Human skin

Plastic surgery skin included reticular dermis to the depth of the fat layer. The upper 200 microns of skin were harvested using a split skin graft knife. Whole skin was digested in RF10 with 1.6mg/ml type IV collagenase for 12-16 hours at 37°C and 5% CO2. Digest was passed repeatedly through a 10ml pipette until no visible material remained. To yield a single cell suspension, digest was passed through a 100-micron filter into a polypropylene sorting tube. Wells were washed twice using cold sort buffer without calcium or magnesium to collect residual and adherent cells.

#### Isolation of leukocytes from Human colon

Normal regions of bowel adjacent to resected colorectal tumours were prepared as previously described, with minor modifications (Bettelli et al. 2006; Geremia et al. 2011). In brief, mucosa was dissected and washed in 1 mM dithiothreitol (DTT) solution for 15 min at room temperature to remove mucus. Specimens were then washed three times in 0.75 mM EDTA to deplete epithelial crypts and were digested for 2h in 0.1 mg/ml collagenase A solution (Roche, UK). For enrichment of mononuclear cells, digests were centrifuged for 30 min at 500g in a four-layer Percoll gradient and collected at the 40%/60% interface.

#### Peripheral blood mononuclear cell isolation

10ml blood from skin donors were collected into EDTA. Density centrifugation with Lymphoprep was performed according to manufacturer instructions. Recovered cells were cryopreserved by pelleting and resuspending in 1ml heat-inactivated fetal calf serum containing 10% DMSO, and storing at −80ºC.

Cryovials were later thawed in water bath, then rapidly being transferred to warmed medium (RPMI 1640 with 100IU/ml penicillin, 10ug/ml streptomycin, 2mM L-glutamine, 10% heat-inactivated fetal calf serum) and filtered through a 100-micron filter.

### Flow cytometry and single-cell RNA sequencing

Mouse and human cell suspensions were stained with the antibodies in Supplementary Table 1 and DAPI. Single cells were sorted in 2μl of Lysis Buffer (1:20 solution of RNase Inhibitor (Clontech, cat. no. 2313A) in 0.2% v/v Triton X-100 (Sigma-Aldrich, cat. no. T9284)) in 96 well plates, spun down and immediately frozen at −80 degrees. Smart-seq2 protocol (Picelli et al. 2014) was largely followed to obtain mRNA libraries from single cells. Oligo-dT primer, dNTPs (ThermoFisher, cat. no. 10319879) and ERCC RNA Spike-In Mix (1:50,000,000 final dilution, Ambion, cat. no. 4456740) were then added. Reverse Transcription and PCR were performed as previously published (Picelli et al. 2014), using 50U of SMARTScribe™ Reverse Transcriptase (Clontech, cat. no. 639538). The cDNA libraries for sequencing were prepared using Nextera XT DNA Sample Preparation Kit (Illumina, cat. no. FC-131-1096), according to the protocol supplied by Fluidigm (PN 100-5950 B1). Libraries from single cells were pooled and purified using AMPure XP beads (Beckman Coulter). Pooled samples were sequenced aiming at an average depth of 1 Million reads/cell, on an Illumina HiSeq 2500 (paired-end 100-bp reads) or Illumina HiSeq 2000 v4 chemistry (paired-end 75-bp reads).

### RNA expression quantification

Gene expression from scRNA-seq data was quantified in Transcripts Per Million (TPM) using Salmon v0.6.0 (Patro et al. 2017), with the parameters --fldMax 150000000 --fldMean 350 --fldSD 250 --numBootstraps 100 --biasCorrect --allowOrphans --useVBOpt. For mouse, the cDNA sequences used contain genes from GRCm38 and sequences from RepBase, as well as ERCC sequences and an EGFP sequence. Since the EGFP RNA is transcribed together with Foxp3, TPM from these two genes were added after quantification to represent Foxp3 expression. For human data quantification, cDNA sequences from GRCh38 and ERCC were used.

### scRNA-seq quality control

After expression quantification, TPM values for each cell were grouped in an expression matrix. ERCC expression levels were separated and TPM were rescaled to total 1 million per cell. Cells were then filtered based on different quality parameters calculated for each dataset (Supplementary Figure 2, Supplementary Table 2) Additionally, the output of TraCeR (Stubbington et al. 2016) was used to remove cells without a detected TCR sequence, as well as iNKT and γ∂ T cells (defined as cells with at least one γ and one ∂ chain detected and no αβ pair). Two mouse steady-state skin Treg from the Mouse Melanoma dataset were also removed from posterior analysis because, even though they had reconstructed TCR and expression of expected T cell genes, they also presented transcripts expected to appear in other cell types, possibly keratinocytes or Langerhans cells (Katayama et al. 2015; Heng, Painter, and Immunological Genome Project Consortium 2008) (Supplementary Figure 16D).

### Dimensionality reduction methods

To obtain an overview of the datasets showing the relationships between cell population clusters, Principal Component Analysis (PCA, Supplementary Figure 3A and 3B) and tSNE were used. tSNE was performed using the Rtsne R package, with the function Rtsne that uses the Barnes-Hutt implementation of the algorithm. For each dataset, a different number of Principal Components (PCs) and values for perplexity were used (Supplementary Table 2). Datasets were treated separately as much as possible to avoid confounding batch effects from experiments performed separately.

### Cell cycle analysis

To assess potential effects of cell cycle in the interpretation of the scRNA-seq datasets collected, Cyclone (Scialdone et al. 2015) (implemented in the scran R package) was used on all datasets. Results were projected on the tSNE (Supplementary Figure 17). As the vast majority of cells was assigned to the default stage (G0/G1 in mouse, S in human), no cell cycle correction was performed.

### Differential expression analysis

Differential Expression (DE) analysis throughout the paper followed the same framework. Two linear models were fitted using the lm function from the base R package to the expression of each gene: a full model containing the information for each population under study (*e.g.* NLT vs non-NLT, Treg vs Tmem, etc.), and a reduced model only including an intercept term. These were then compared using a likelihood-ratio test (lmtest R package), and a q-value was calculated to correct for multiple testing. For all tests between tissues, genes with a q-value<=0.01 and a log2 fold-change>=1 were considered differentially expressed. For tests between cell types within a tissue, only genes detected as markers for that tissue were considered, and a q-value threshold of 0.05 was used. When calculating differentially expressed genes in human skin, or between skin and blood, the donor effect was included as part of the null model so as not to impact the test. In the human and mouse comparison, human NLTs were compared to blood and mouse NLTs were compared to spleen only.

### Differential co-expression analysis

Cell identity is defined not only by expression of specific genes, but also interactions between them. To assess which interactions define Treg and Tmem identity in NLTs or Treg identity between NLTs, we used the DGCA package (McKenzie et al. 2016), which relies on the Fisher z-score transformation of the Spearman correlation values to perform a differential correlation test on gene pairs. For these tests, only genes identified as NLT markers and expressed in more than 5 cells in both conditions tested were used. We only kept differentially correlating genes where at least one population had a correlation coefficient greater than 0.25 (meaning co-expression of the gene pair in the tested condition).

### Obtaining a migration latent variable for steady-state Tregs

The large dimensionality of single-cell RNA-seq data has been used before to gain insights on time-dependent events (Trapnell et al. 2014; Lönnberg et al. 2017) by applying methods for pseudotime inference. Although it is impossible to follow one cell through the complete process, these methods can order single-cell data into a continuous dimension, using the discrete samples as snapshots containing a multitude of intermediate states.

Immune cells are expected to migrate via blood or lymph. We assumed that this effect would be reflected as a gradual single-cell expression phenotype, which could be captured as a latent variable of the data. To achieve this, we used Bayesian Gaussian Process Latent Variable Modelling (BGPLVM) (Michalis K Titsias 2010), implemented in the python package GPy (https://github.com/SheffieldML/GPy) as “GPy.models.BayesianGPLVM”, which was already used before for dimensionality reduction in scRNA-seq data to model Th1-Tfh differentiation (Lönnberg et al. 2017). BGPLVM was used asking for 6 latent variables, and the two most significant by Automatic Relevance Determination (ARD, Supplementary Figure 10) were plotted. We excluded spleen cells from this analysis to avoid making assumptions about the migration direction between the three tissues, and focus on general LN to NLT trafficking. This analysis was performed separately in Treg and Tmem, to avoid a cell type-specific confounding effect (Figure 4A and 4D, Supplementary Figure 13). Indeed, while skin Tmems appear to have latent effects similar to those present in Treg (Supplementary Figure 13F), the same can not be said for Tmems from the mouse colon dataset, where only a division by tissue can be observed (Supplementary Figure 13D). This might be due to further specialization of colonic Tmems, which we observed when searching for subpopulations (Supplementary Figure 5D-F).

We observe that gene correlations with colon LV1 and skin LV0 are correlated (Supplementary Figure 10 C), and a high proportion of those genes are associated with NLT identity (Supplementary Figure 10D). We see that the effect from these LVs is also present in each tissue individually, unlike what is observed for colon LV0 or skin LV1 (Supplementary Figure 11). Thus, we identified colon LV1 and skin LV0 as representing tissue adaptation in each dataset. Lastly, intervals for each tissue (Figure 4, Supplementary Figure 14) were defined in each 11 cell interval what tissue was more represented.

### Identifying a common tissue migration trajectory in control and melanoma

Similarly to the steady state, migration from the LN to the skin with a melanoma challenge is also expected. A common between-tissue Treg migration trajectory in control and melanoma conditions was obtained using Manifold Relevance Determination (Andreas Damianou, Carl Ek, Michalis Titsias, Neil Lawrence 2012) (MRD). MRD works by having an underlying BGPLVM model whose dimensions can be shared or private between sections of the data. The importance of each section in each latent variable is shown in the ARD plot (Supplementary Figure 18). Having the prior knowledge that a cell-cycle effect is present in the data (Supplementary Figure 17) and with the goal of obtaining a latent variable explaining tissue recruitment in both conditions, the melanoma dataset was divided into three sections for input: one with the expression in all cell-cycle associated genes, one with marker genes for any tissue, and one with the remaining genes. The model was run allowing for 12 latent variables as output, and the one highly influenced by tissue-specific genes but not cell-cycle or other genes was used as a migration trajectory for both conditions. The effects captured by these latent variables can be observed in BGPLVM projections for the individual conditions (Supplementary Figures 19A-D), and the latent variable with the largest tissue identity is also the most correlated to the LV used in the skin steady-state (Supplementary Figures 19E). However, the cell-cycle associated latent variable also includes tissue-related effects, which can be explained by the expansion of Treg just before leaving the LN and upon arriving in the skin.

### Switch-like genes in the migration latent variable

Gene expression changes in a continuous trajectory can be interpreted as a series of switch-like events. These can be modeled using a sigmoid curve, described by the following equation:

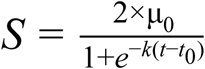

where μ_0_ is the mean expression between the sigmoid “on” and “off” states, *t*_0_ is the point in which the switch in expression happens, and *k* defines the sigmoid inclination and can be interpreted as the activation strength. Parameter *k* will additionally inform on the direction of the switch (activation or inhibition) from its signal.

The R package switchde (Campbell and Yau 2017) was used to model gene expression as a sigmoid in the inferred migration trajectories, using the appropriate latent variable as pseudotime. In the steady-state datasets, switchde was applied for genes detected as Treg markers in either dataset (Treg vs Tmem differential expression regardless of tissue), tissue markers for each tissue in each dataset, or genes with an absolute correlation greater than 0.25 with the latent variable chosen. For the melanoma dataset, genes expressed in at least 5 cells in both conditions were tested. Only genes with a q-value <=0.05 and that had a *t*_0_ within the LV range were kept for further interpretation.

### Detection of expanded clonotypes

T cell receptor (TCR) sequences were reconstructed from single cell RNA-seq data and used to infer clonality using TraCeR (Stubbington et al. 2016). We used TraCeR with the parameters --loci A B D G, --max_junc_len 120 to allow reconstruction of TCRα, TCRβ, TCR∂ and TCRγ chains in each cell and to permit TCRγ chains with long CDR3 regions.

### GO Term enrichment

To test for enriched GO Biological Processes or KEGG Pathways in gene sets, the gprofiler R package (Reimand et al. 2016) was used, with the option of moderate hierarchical filtering enabled.

### Clustering analysis

To search for subpopulations in the NLT obtained Treg and Tmem cells, we used the consensus clustering algorithm implemented in the SC3 R package (Kiselev et al. 2017). We looked for 2 to 4 clusters, to allow us to capture any possible subpopulations, but also be able to exclude cases where small spurious subpopulations are identified. For each tissue/cell combination, we chose to compare the largest subpopulations detected that also were the farthest apart in tSNE space. Differential expression between clusters was performed on all expressed genes in them.

It is worth noting that absence of evidence for the existence of subpopulations is not evidence of their absence. Indeed, it has previously been shown that the number of cells in a scRNA-seq experiment is more important for subpopulation detection than the number of detected genes (Shekhar et al. 2016). Therefore, we underline that the presented datasets are mostly useful for a deep characterization of the sorted populations.

## Author’s contributions

RJM, AC, AH, TK, FP and SAT conceived the project and designed steady-state experiments. RJM and AR designed the melanoma challenge experiments.

AC and AH collected cells for steady-state mouse and human colonic datasets. LJ collected cells for human skin dataset. AR induced the melanoma challenge. RJM and AR collected cells for melanoma dataset. RJM performed scRNA-seq. TG, RJM and SAT planned the data analyses. TG and RJM analyzed the data. IL performed the TraCeR analysis. RJM, TG, AC, SAT wrote the manuscript. SAT, FP, MH and JS supervised the work.

## Acknowledgements

We thank V. Proserpio, M. Stubbington, T. Hagai, X. Chen, F.V. Braga, V. Svensson, J. Henriksson for helpful discussions and advice. We thank T. Hagai, R. Vento-Tormo, K. Meyer, J Park for critical reading of the manuscript.

We thank the WTSI Single-cell Genomic Core Facility, WTSI Sequencing Facility, WTSI Flow Cytometry facilities, as well as the Kennedy Institute Flow Cytometry Facility, for expert technical advice and assistance.

RJM was supported by a PhD Fellowship from the Fundação para a Ciência e Tecnologia, Portugal (SFRH/BD/51950/2012), TG by the European Union’s H2020 research and innovation programme “ENLIGHT-TEN” under the Marie Sklodowska-Curie grant agreement 675395. This project was supported by ERC grants ThDEFINE and ThSWITCH. ANH was supported by an EMBO long-term fellowship (ALTF 1161-2012) and a Marie Curie fellowship (PIEF-GA-2012-330621).

## Data Accessibility

scRNA-seq data for this project has been deposited in ArrayExpress under the accession number E-MTAB-6072.

